# Different serotonergic neurons regulate appetite for sucrose and hunger for proteins

**DOI:** 10.1101/2024.10.30.621076

**Authors:** Katharina Dorn, Magdalena Gompert, Jianzheng He, Henrike Scholz

**Affiliations:** Department of Biology, Institute for Zoology, University zu Köln, 50674 Köln, Germany

**Keywords:** serotonin, hunger, appetite, insulin receptor, serotonin transporter, carbohydrates, proteins

## Abstract

For the organism it is important to replenish internal energy storages selectively and selective appetite for nutrients might uncover internal energy requirements. How is the selective uptake of a specific nutrient regulated? Here we show that in *Drosophila melanogaster* different sets of serotonergic neurons regulate appetite for sucrose and hunger for proteins. Increased neuronal activity in specific subsets of serotonergic neurons and interfering with serotonin reuptake using a mutated serotonin transporter reduced the appetite for sucrose selectively, but not the hunger for proteins. The insulin receptor together with the serotonin transporter regulates the selective sucrose appetite. We provide evidence that the cellular location of the serotonin transporter depends on the insulin receptor. This mechanism might allow optimizing nutrient intake and in turn might prevent overconsumption by repressing appetite for sucrose. Given the conserved nature of the molecules involved it is likely that the mechanism is conserved in higher organisms.

## INTRODUCTION

Food intake must be regulated depending on the organism’s internal energy requirements and the availability of external food sources. The lack of internal energy manifests itself by the feeling of hunger. Hunger leads to pronounced foraging behavior and increases the sensitivity to external stimuli to ensure the detection of a suitable food source (Ko et al., 2015). Appetite, on the other hand, is defined as the desire to feed despite the lack of hunger and can optimize the intake of certain nutrients. Loss of control over appetite-driven food intake in humans often leads to overeating and metabolic disorders such as obesity and diabetes (Berthoud & Morrison, 2008). In vertebrates and invertebrates, the neurotransmitter serotonin is involved in the regulation of appetite and hunger (Tierney, 2020). However, it remains unclear how serotonin regulates selective appetite for certain nutrients.

In vertebrates, increased serotonin levels are associated with the repression of appetite (Voigt & Fink, 2015; Yabut et al., 2019). In adult *Drosophila melanogaster*, the role of serotonin in food intake is diverse and serotonin regulates different aspects of feeding (Albin et al., 2015; Pooryasin & Fiala, 2015; Ro et al., 2016; Yao & Scott, 2022). For example, in satiated flies, the activation of subsets of serotonergic neurons results in increased food intake (Albin et al., 2015), whereas in hungry flies the activation of a similar and broader set of serotonergic neurons led to a reduction in food intake (Pooryasin & Fiala, 2015). Preference for a high-protein diet after mild starvation depends on serotonin levels (Ro et al., 2016). Different serotonergic neurons respond with activity to sucrose or bitter taste. The serotonergic neurons that respond to sucrose activate insulin secretion, while the neurons that respond to bitterness regulate gastric motility (Yao & Scott, 2022).

The presence of serotonin in the synaptic cleft can be prolonged by blocking the reuptake of serotonin via the serotonin transporter into the presynaptic neuron. In rats, serotonin reuptake inhibitors reduce food intake (Halford et al., 2007; Simansky & Vaidya, 1990). In humans, serotonin reuptake inhibitors have been used as anorexic drug to reduce food intake and weight (Hainer et al., 2006; Voigt & Fink, 2015). The mechanism appears to be conserved between flies and mammals, as flies lacking the *Drosophila serotonin transporter* (SerT) also show a reduced food intake (Knapp et al., 2022).

Regulation of food intake affects the amount of nutrients available to produce or store energy on cellular level. At the cellular level, insulin receptors regulate the uptake of glucose into the cells (Saltiel & Kahn, 2001). In vertebrates and invertebrates such as *Drosophila melanogaster*, the signal transduction cascades acting downstream of the insulin receptors are well conserved (Chatterjee & Perrimon, 2021; Inoue et al., 2018). In contrast to insulin in vertebrates, several insulin-like peptides have been identified in *Drosophila* that could interact with the insulin receptor (Chowanski et al., 2021). In *Drosophila*, insulin-like peptides (Dilp) regulate food intake (Semaniuk et al., 2021). Several *Dilp* mutants consume too much sucrose or protein-enriched food (Semaniuk et al., 2021; Semaniuk et al., 2018), suggesting that the activation of insulin receptor signaling via different Dilps might serve as satiety signal. Under starvation, this might change, as Dilps promote food intake (Sudhakar et al., 2020). Alternatively, different downstream targets of Dilp signaling can regulate different aspects of selective food intake in context of different internal energy demands.

To better understand how internal energy resources are replenished, we investigated how serotonergic neurons regulate nutrient-specific food intake and analyzed the role of serotonin transporter and insulin receptor function in the regulation of this specificity. Using behavioral experiments and neuroanatomical analysis, we identified a group of serotonergic neurons that specifically regulate the appetite for sucrose, but not high-protein diet. Increased serotonin signaling reduced sucrose appetite similar to a blocked or activated insulin receptor function. The reduction in sucrose appetite could be due to two different mechanisms. One mechanism is due to changes in the morphology of serotonergic neurons, that can reduce sucrose intake. The second mechanism is due to dislocation of the serotonin transporter within the neurons. Using genetic epistasis experiments, we provide evidence that the serotonin transporter acts downstream of insulin receptor in regulating appetite of sucrose. Reduced appetite for sucrose could in turn affect other appetitive behaviors, such as learning and memory of food reward. Together, our results reveal a mechanism by which insulin receptor signaling regulate the appetite for a specific nutrient. This mechanism enables nutrient-specific food intake required to selectively replenish internal energy stores.

## RESULTS

General activation of serotonergic neurons suppresses food intake (Pooryasin & Fiala, 2015). However, in satiated flies, activation of serotonergic neurons triggers food intake (Albin et al., 2015). This raises the question of how the serotonergic neurotransmitter system does both: suppressing and enhancing food intake.

### Different sets of serotonergic neurons regulate appetite or hunger for certain nutrients

The difference in food intake might depend on the set of serotonergic neurons that regulate the intake, the internal energy state of the fly and/or the food source. Therefore, we wanted to investigate the function of different groups of serotonergic neurons in flies with different energy states during the food intake on different diets (Figure 1).

**Figure 1.**
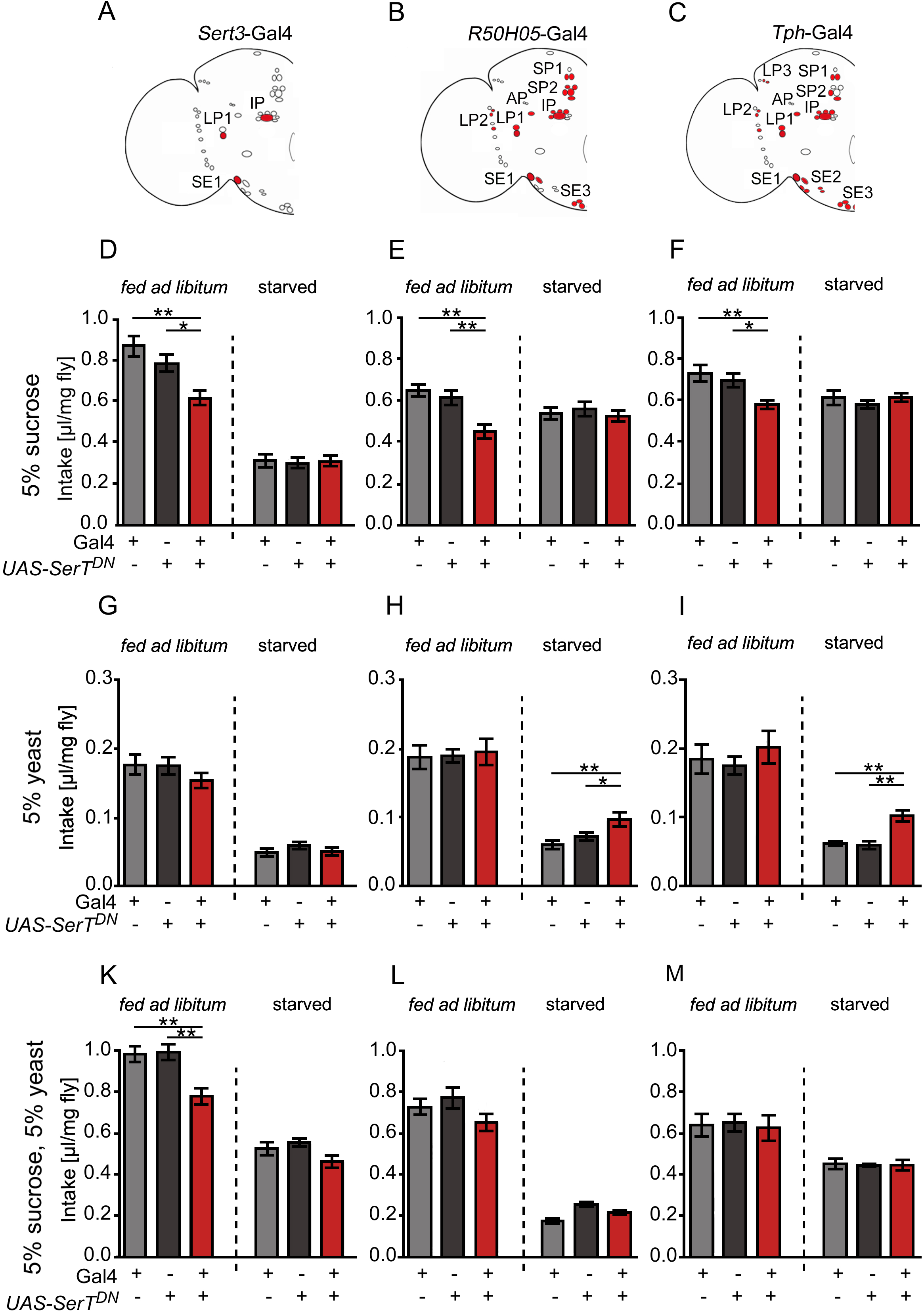
Serotonergic neurons regulate sucrose appetite and protein hunger. **A** to **C**, Schematics of serotonergic neurons targeted by different Gal4 drivers; serotonergic neurons are marked in red. **D** to **F**, Expression of SerT^DN^ under the control of the *Sert*-Gal4, *R50H05*-Gal4 or *Tph*-Gal4 drivers reduced sucrose intake in flies *fed ad libitum* but not starved flies. **G** to **I**, Hunger-induced protein intake is increased, when SerT^DN^ is expressed in broad subsets of serotonergic neurons. **K** to **L**, Appetite-induced intake of a mixed diet consisting of 5% sucrose and 5% yeast is reduced only in flies in which SerT^DN^ is expressed under the control of the *Sert3*-Gal4 driver. Data represents mean ± SEM (n = 16–37). For raw data see Supplementary Table S1. \**P* < 0.05; \*\**P* < 0.01; one-way ANOVA, with Tukey post-hoc test.

To specifically alter serotonergic function in serotonergic neurons, we employed the expression of a non-functional serotonin transporter with mutated serotonin binding sites under the control of the Gal4/*UAS* system that competes with the endogenous serotonin transporter for function. The serotonin transporter exists as a multimer that takes up serotonin from the synaptic cleft after release and thereby terminates transmission between pre-and post-synapsis (Cooper et al., 2019; Rudnick & Sandtner, 2019). The expression of the mutated SerT results in reduced presynaptic serotonin and functions as a dominant-negative serotonin transporter (*UAS*-*SerT^DN^*; (Xu et al., 2016; He et al., 2020). To target different sets of serotonergic neurons, we used three different serotonergic Gal4 drivers (Figure 1A to Figure 1C). The *Sert3-* Gal4 driver targets three serotonergic neurons per brain hemisphere, including one SE1 neuron, one IP neuron, and one LP1 neuron (Xu et al., 2016). The *R50H05-*Gal4 and the *Tph*-Gal4 drivers target approximately 54% of serotonergic neurons, which comprise partially overlapping sets of serotonergic neurons (Jenett et al., 2012; Park et al., 2006; Xu et al., 2016). To investigate the influence of the internal energy state, we analyzed food intake of starved flies to determine hunger-induced food intake or of *ad libitum*-fed flies to determine appetite-induced food intake. We used 5% sucrose, 5% yeast or 5% sucrose with 5% yeast as food source and determined food intake using a Capillary Feeder assay (CaFe assay; Ja et al., 2007; Figure 1).

Expression of the *UAS*-*SerT^DN^* under the control of *Sert3-*Gal4*, R50H05-*Gal4, and *Tph*-Gal4 reduced appetite for sucrose but did not alter appetite for yeast (Figure 1D to Figure 1F; Figure 1G to Figure 1I). The same sets of serotonergic neurons did not regulate sucrose hunger (Figure 1D to Figure 1F). Expression of the *UAS-SerT^DN^* in a set of serotonergic neurons under the control of the *R50H05-*Gal4 and the *Tph*- Gal4 drivers increased the yeast hunger (Figure 1H to Figure 1I). When comparing the different sets of serotonergic neurons targeted by the different drivers, the reduced appetite for sucrose can be attributed to three serotonergic neurons targeted by the *Sert3*-Gal4 driver, while protein hunger is regulated by additional neurons of clusters SE3, IP or LP1 or serotonergic neurons of the clusters SE3, SP1, SP2, LP2 or AP.

Next, we investigated how information from hunger-sensing and appetite sensing-neurons jointly influences food intake of food containing sucrose and yeast. We therefore gave starved flies a mixed diet of 5% sucrose and 5% yeast or gave fed flies *ad libitum* excess to the food source (Figure 1K to Figure 1M). Although expression of SerT^DN^ under the control of the *Sert3*-Gal4 driver still resulted in reduced appetite for the food mixture, it had no effect on hunger-induced food intake (Figure 1K). Expression of the SerT^DN^ under the control of the *R50H05-*Gal4 and *Tph*-Gal4 did not alter appetite or hunger for the mixed food diet, suggesting that serotonergic neurons exist that counteract the reduced appetite for sucrose when a balanced diet is provided. Similarly, hunger for proteins can be suppressed by a diet high in proteins and sucrose.

In summary, appetite for sucrose and hunger for a high-protein diet are regulated by different sets of serotonergic neurons. In the presence of a food source containing sucrose and proteins, the information from different serotonergic neurons is processed together and information of sucrose appetite and protein hunger is offset against each other.

### Activation of serotonergic neurons reduce appetite for sucrose or hunger for proteins

Expression of a mutated serotonin transporter during development could lead to compensatory effects in the brain. To analyze whether activation of serotonergic neurons directly regulate sucrose intake, we activated the set of serotonergic neurons using optogenetics in adult flies and analyzed food intake. To do so, we combined the CaFe assay with a light source that allowed to activate the light-sensitive Channelrhodopsin-2 (ChR2) under the control of the *Sert3*-Gal4 or *Tph*-Gal4 driver (Figure 2). Blue light irradiation causes a conformational change of ChR2 in the presence of all-trans retinal (ATR), which leads to influx of cations and depolarization of neurons, thereby increasing neurotransmitter release (Nagel et al., 2003). We analyzed the food intake of hungry flies or flies fed *ad libitum* (Figure 2B to Figure 2D). As control group, we used flies that were not fed with ATR. In addition, we controlled for the effect of ATR on the ChR2 transgene by examining food intake of flies carrying one copy of the *UAS-ChR* transgene.

**Figure 2.**
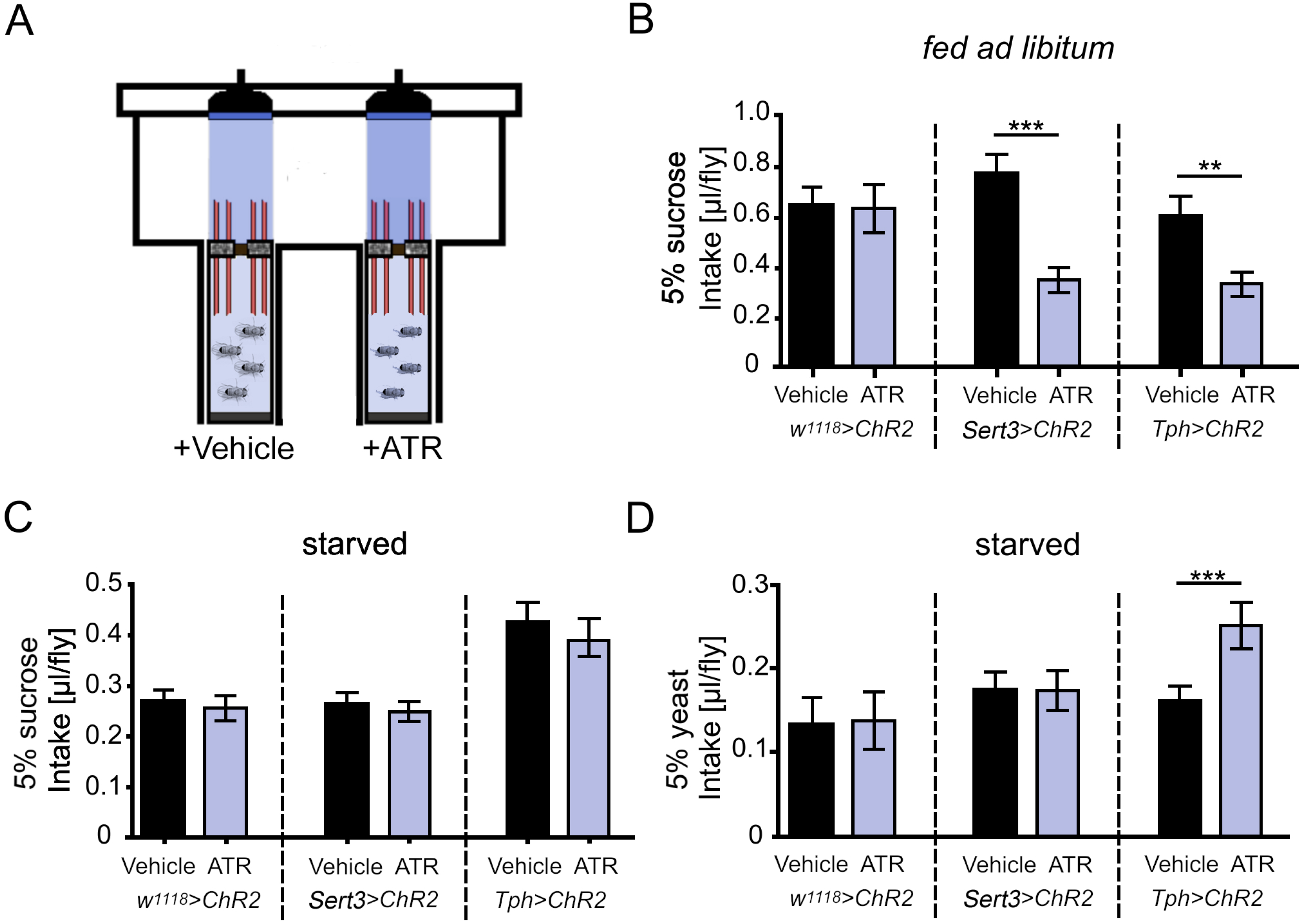
Activation of serotonergic neurons decreases appetite for sucrose and hunger for proteins. **A**, The CaFe-assays were combined with blue diodes with a light intensity of 1800 lx and a flicker frequency of 2 s 40 Hz, 16 s 8 Hz and 2 s of no light. **B**, Activation of serotonergic neurons decreased appetite-induced sucrose intake- **C**, Activation does not affect hunger-induced sucrose intake. **D**, Hunger-induced protein intake increases with activation of a broad subsets of serotonergic neurons. Data are mean ± SEM (n = 16-21). For raw data see Supplementary Table S1. Unpaired Student’s t-test for comparison between the groups: \*\**P* < 0.01, \*\*\**P* < 0.001.

Activation of serotonergic neurons under the control of the *Sert3*-Gal4 or *Tph*- Gal4 driver reduced appetite-induced sucrose intake (Figure 2B), but not hunger-induced sucrose intake (Figure 2C). This supports our results showing that the serotoninergic neurons function as negative regulator of sucrose appetite. Consistent with the idea that a different set of serotonergic neurons regulates protein hunger, we observed activation of neurons targeted by the*Tph-*Gal4 driver increased hunger-induced protein intake, but not activation of the serotonergic neurons targeted by the *Sert3*-Gal4 driver (Figure 2D). Thus, different sets of serotonergic neurons regulate sucrose appetite and protein hunger.

### Reduced appetite correlates with reduced appetitive short-term memory

To clarify whether a reduced appetite for sucrose is also associated with a reduction in the rewarding properties of sucrose, we performed classical olfactory conditioning experiments with sucrose as a reinforcer. To do so, we used the Tully- Quinn paradigm and analyzed whether flies expressing the *UAS*-*SerT^DN^* under the control of the *Sert3-*Gal4 driver form a positive association between an odorant and sucrose within 2 min (Figure 3).

**Figure 3.**
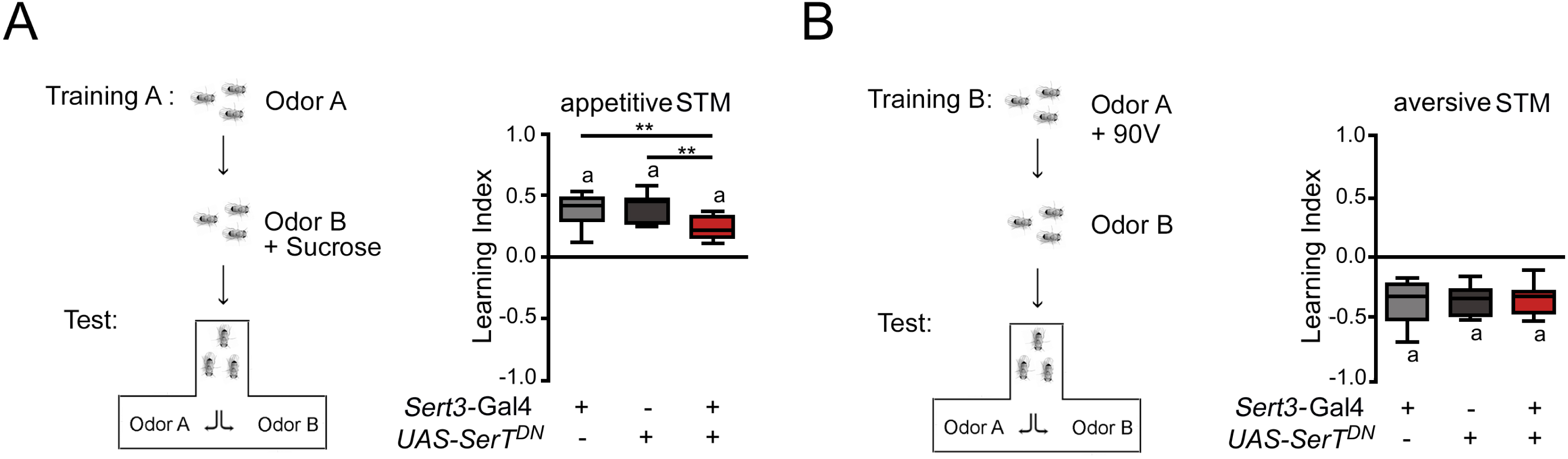
Influence of reduced sucrose appetite on appetitive and aversive STM. **A,** To assess appetitive short-term learning and memory, flies were first exposed to one odorant paired with a reward of 2 M sucrose followed by a second odorant during training. **B,** To asses aversive learning and memory, flies were exposed to the odor paired with an electric shock of 90V for 1 min followed by an exposure to a second odor (US) of 1 min. In the test situation 3 min after the training, they choose for 2 min between both odorants **A**, Expression of *UAS*-*SerT^DN^* under the control of *Sert3*-Gal4 significantly reduced the olfactory appetitive STM. **B**, Aversive STM is not affected in flies with expression *UAS*-*SerT^DN^* under the control of *Sert3*-Gal4. The letter “a” indicates a significant difference from random choice (one-sample t-test (*P* < 0.05)). N = 12. For raw data see Supplementary Table S1. Statistic differences between groups were determined by a One-way ANOVA followed by post-hoc Tukey HSD test; ** *P* < 0.01.

To assess appetitive short-term learning and memory, flies were exposed to the 3-octanol for 2 min and then to the second odorant 4-methylcyclohexanol with a reward of 2 M sucrose (Schwaerzel et al., 2003; Tully & Quinn, 1985). After training, the flies were given the choice between the reinforced and non-reinforced odorant for 2 min (Figure 3A). A second independent group of flies were trained in a reciprocal manner. The learning indices for both reinforced odorants were used to determine the average learning index (LI). Flies expressing the SerT^DN^ under the control of the *Sert3-Gal4* driver showed reduced appetitive short-term learning and memory (Figure 3B).

To address whether general mechanisms underlying short-term learning and memory are impaired, we performed similar experiments using a 90 V electric shock as a negative reinforcer (Figure 3B). Here, we observed no differences between the experimental group and the corresponding controls (Figure 3B). The reduction in appetitive STM was not due to differences in sensory acuity, as the flies sensed the odorants and the reinforcers, sucrose and the electric shock, in a similar way (Table 1). Thus, the reduced appetite of sucrose correlates with reduced reward properties of sucrose.

**Table 1.** Sensory acuity tests in serotonergic neurons targeted by *Sert3*-Gal4. Mean ± s.e.m is shown. No significant differences between control and experimental groups were detected using ANOVA Turkey-Cramer post hoc analysis (*P* < 0.05). The letter **a** indicates significant differences from random choice as determined by One-sample sign test (*P* < 0.05). N = 12 - 18 for each test except sucrose sensitivity (N = 20).

| Genotype | Sucrose reactivity | Shock reactivity | Odor avoidance |  | Balance |
| --- | --- | --- | --- | --- | --- |
|  |  |  | 3-Oct (1:80) | MCH (1:100) |  |
|  |  |  | Starved | Starved |  |
| *Sert3-Gal4* | 0.72±0.08a | -0.87±0.04a | -0.38±0.06a | -<br>0.32±0.05a | 0.03±0.06 |
| *UAS-SertDN* | 0.66±0.07a | -0.85±0.03a | -0.37±0.05a | -<br>0.32±0.05a | 0.07±0.06 |
| Experimental | 0.66±0.07a | -0.86±0.04a | -0.35±0.05a | -<br>0.26±0.05a | 0.09±0.07 |

### The Insulin-like receptor regulates appetite for sucrose

At the cellular level, the insulin-like receptor in *Drosophila* regulates glucose uptake, a metabolite of sucrose (Chatterjee & Perrimon, 2021a). Expression of the activated insulin-like receptor reflects high internal glucose levels and should lead to suppression of food intake. To assess whether the insulin-like receptor regulates the appetite for sucrose in serotonergic neurons, we expressed the constitutively active InR (InR^CA^) under the control of the *Sert3*-Gal4 driver and analyzed food intake (Figure 4). Expression of the active insulin-like receptor reduced appetite for sucrose, but not hunger for sucrose (Figure 4A). Similarly, the appetite for sucrose and yeast-containing food was reduced, but not the hunger for the mixed diet. Appetite for yeast or hunger for sucrose remained unchanged (Figure 4A).

**Figure 4.**
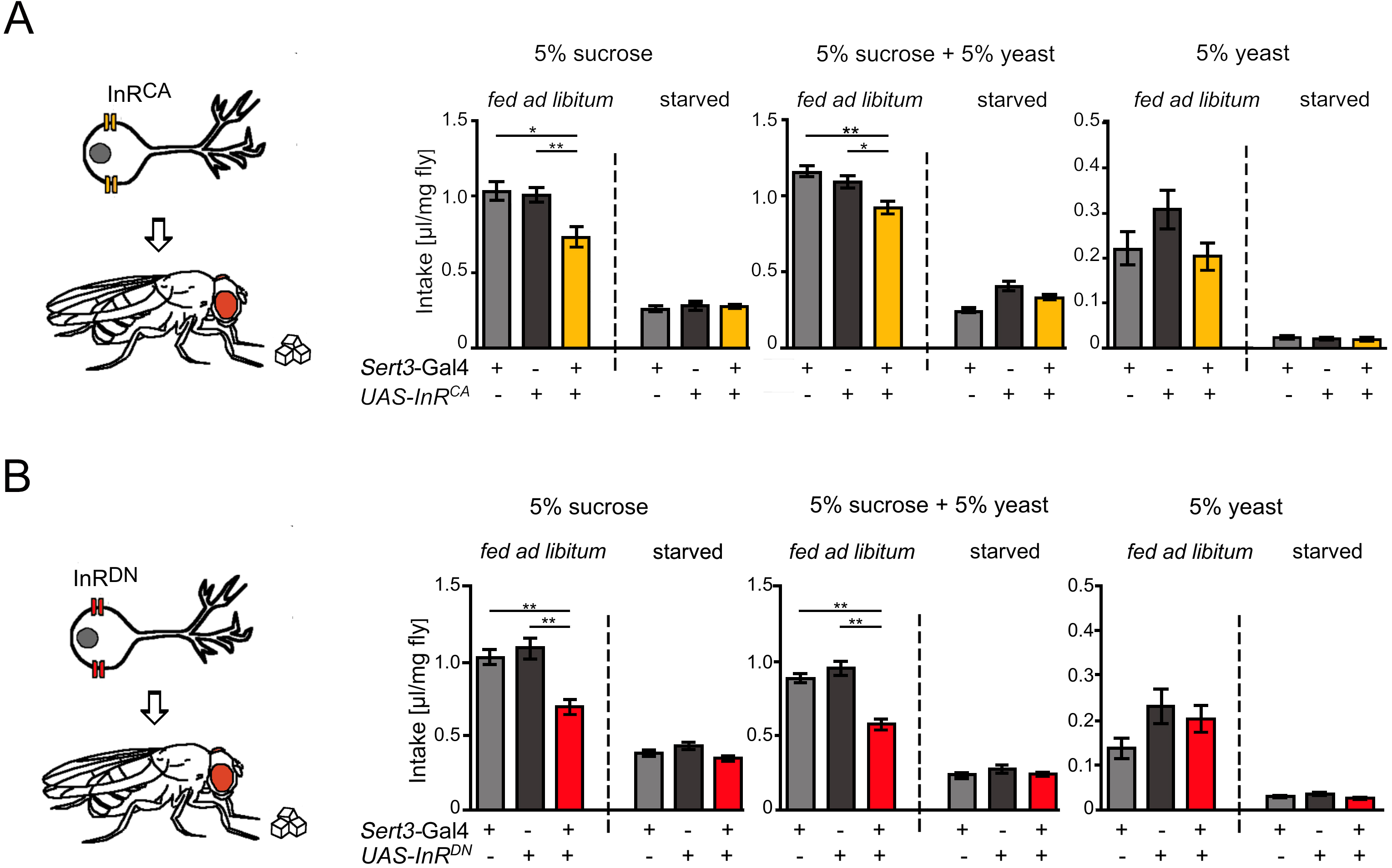
The insulin-like receptor regulates appetite-induced sucrose intake. **A**, Flies expressing InR^CA^ under the control of the *Ser3*-Gal4 driver reduced their food intake when fed ad libitum with sucrose or sucrose and yeast, but not with yeast only. The food intake of hungry flies was not affected. **B**, Expression of InR^DN^ under the control of the *Ser3*-Gal4 driver only reduced food intake in flies that were fed either sucrose or sucrose and yeast ad libitum. The data are mean food intake ± SEM (n = 12-36); for raw data see Supplementary Table S1. \**P* < 0.05, ** *P* < 0.01 based on ANOVA Tukey HSD test.

We next investigated whether blocking insulin-like receptor function would increase appetite-induced food intake (Figure 4B). Blockade of InR should reflect a reduced internal glucose level or a starvation state of the animal. Expression of a dominant negative InR (InR^DN^; Demontis & Perrimon, 2009) under the control of the *Sert3*-Gal4 driver reduced appetite-induced intake of sucrose or a mixture of sucrose and yeast, but did not alter yeast intake. Hunger-induced food intake was not affected, regardless of the type of food. We again observe the selective effect of appetite-driven sucrose intake, but surprisingly, activation and inactivation of the InR led to similar effects.

### The Insulin-like receptor influences SerT expression

Changes in InR signaling or blocking SerT function decreased appetite for sucrose, suggesting that both molecules act in a similar process. To investigate whether the insulin-like receptor affects SerT function, we wanted to analyze whether altered InR signaling alters the expression of SerT (Figure 5). We focused our analysis on neurons within the fan-shaped body (fb) of the brain, since they have distinct morphological domains with clearly visible synaptic regions (Hanesch et al., 1989). First, we compared the expression of the serotonin transporter with serotonin expression (Figure 5A). In line with previous result, there are regional differences in the strength of expression in other structures of the brain, but above all SerT und serotonin expression overlaps (Giang et al., 2011; Figure 5A′′). For example, both occur overlapping in the typical glomerular structure of layer 4 of the fan-shaped body. The *Tph*-Gal4 line drives UAS-mCD8::GFP expression in the serotonergic ExR3 neuron, that innervates the dorsal part of the fan-shaped body and contributes to the glomerular structure in layer 4 (Omoto et al., 2018; Figure 5B).

**Figure 5.**
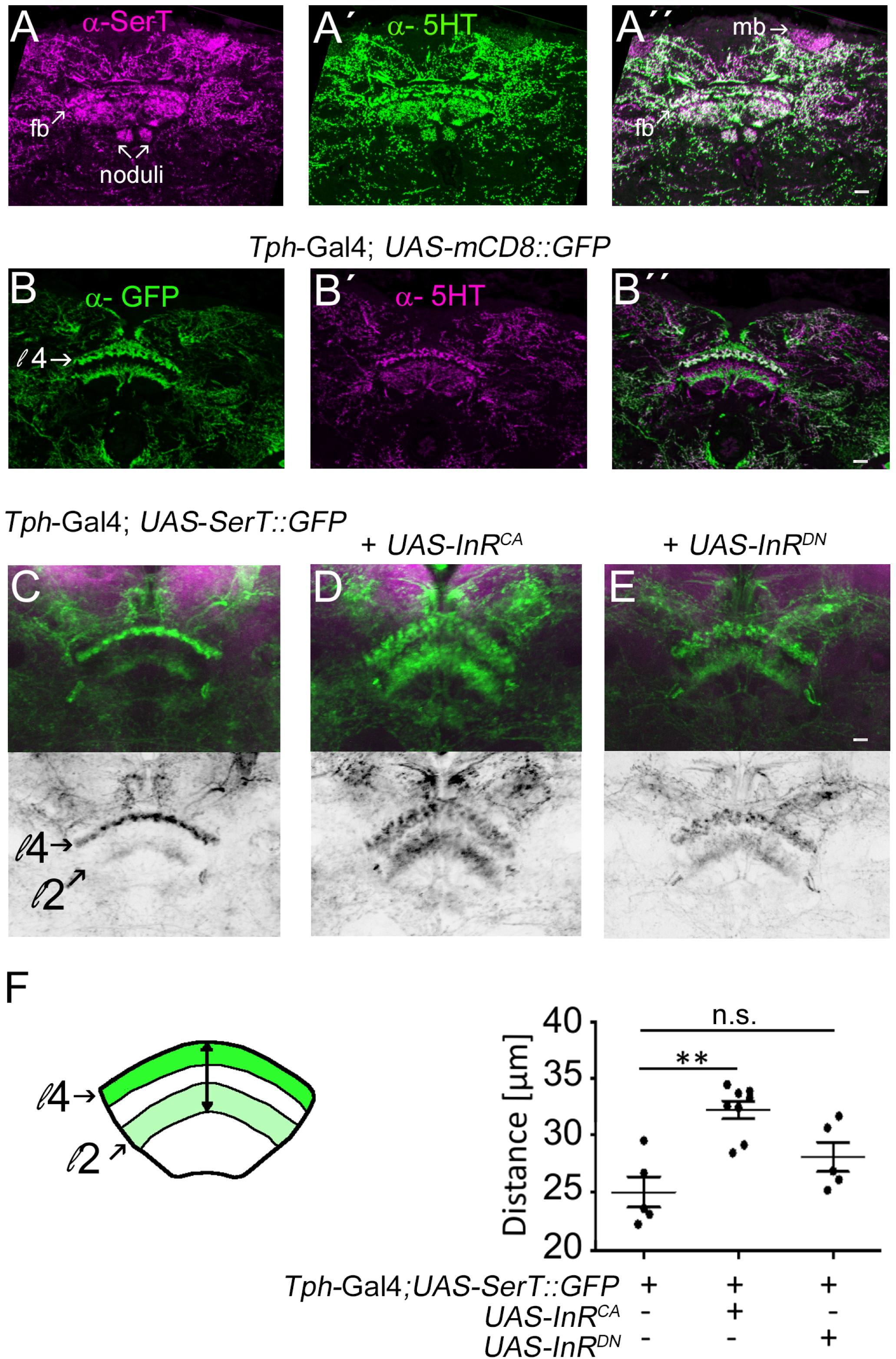
Insulin receptor function affects SerT expression. **A** to **A′′,** Expression of SerT (in magenta), expression of serotonin (in green) and overlap (in white) in 10µm thick brain section showing the fans-shaped body (fb). **B** to **B′′**, The *Tph*-Gal4 driver targets the ExR3 neuron that innervates the fb and contributes to the layer 4 (l4) of the fb in the whole mount brain of a control fly (GFP expression in green and serotonin expression in magenta). **C** to **E**, The GFP signal (in green) of the *UAS-SerT::GFP* transgene under the control of the Tph-Gal4 driver changes the localization and intensity depending on **D**, InR^CA^, or **E**, InR^DN^ , expression. To quantify the expression area of SerT::GFP, the width of layer 2 and layer 4 was measured. The nc82 antigen was used to label neurophils (in magenta) in the whole mount brain. l2: layer 2; l4: layer 4. N = 5 – 8 brains. The data were compared using ANOVA Tukey HSD test. For raw data see Supplementary Table S1. ** *P* < 0.01.

To analyze the SerT expression in response to altered InR function, we used the *UAS-SerT::GFP* transgene (Figure 5C to Figure 5E). The GFP expression is clearly visible in densely packed glomerular structures in layer 4 and lightly in an additional layer (Figure 5C). Co-expression of InR^CA^ results in more intense GFP expression in layer 4 and loosely packed glomerular structures in addition to more intense expression in a second layer mainly containing projections (Figure 5D). Co-expression of InR^DN^ resulted in more punctate GFP expression in layer 4 appearing as loosely packed glomerular structures. The expression of the activated InR broadened the expression of the SerT::GFP as determined by measuring the width between the two layers which was not the case when inactivated InR was expressed (Figure 5E). In summary, the expression of the activated and inactivated InR alters the expression of SerT, but not necessarily in the same way.

### The Insulin-like receptor regulates neuron morphology and SerT localization

To analyze the connection between InR signal and SerT at the level of individual neurons, we next focused on the serotonergic SE1 neuron targeted by *Sert3-*Gal4 driver (Figure 6A). The soma of the SE1 neuron is located in the subesophageal zone (SEZ). In the SEZ, the projection of the SE1 neuron divides into three branches, two of which span the SEZ and the third projects towards the ventral nerve cord. To visualize the morphology of the SE1 neuron, a membrane-bound RFP was expressed under the control of the *Sert3-*Gal4 driver and SerT expression was monitored using a *UAS-SerT::GFP* transgene (Figure 6 B to Figure 6 E). Expression of activated InR^CA^ did not alter the gross morphology of the branching pattern, but reduced SerT::GFP expression in the region of the branches and synaptic varicosities within the SEZ (Figure 6C). In contrast, expression of InR^DN^ leads to a reduction in branching. SerT::GFP expression was mainly found in the soma and the projection targeting into the ventral nerve cord (Figure 6C). This suggests that the reduced appetite for sucrose observed by expression of the activated insulin receptor or the blocked receptor is due to two different effects. Consistent with a function of the InR in axonal growth (Li et al., 2013), blocking insulin receptor function leads to a reduction in axon branching. In contrast, activation of the InR receptor leads to alteration of the intracellular localization of the serotonin transporter.

**Figure 6.**
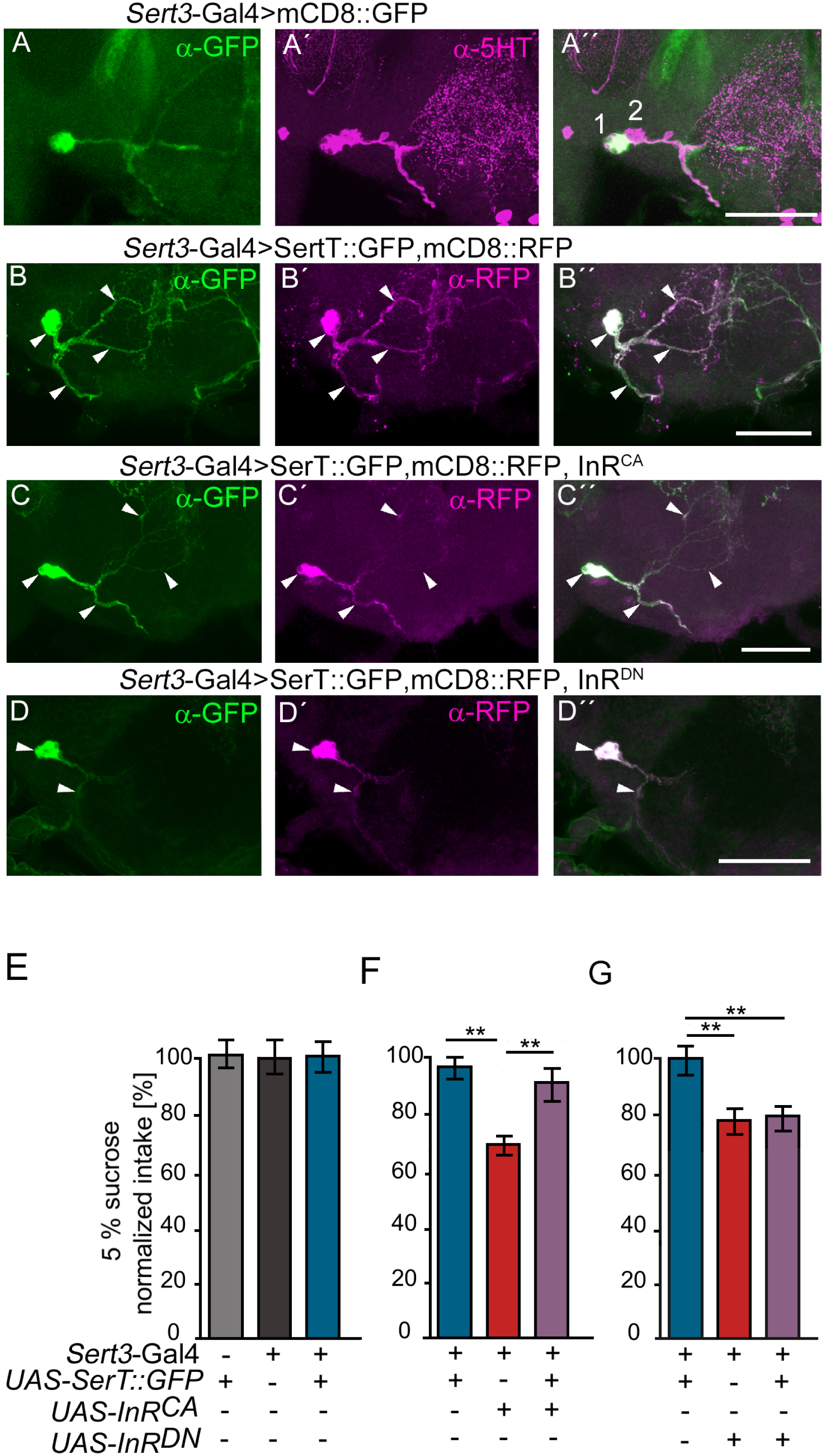
Insulin receptor function regulates neuron morphology and localization of SerT expression. **A**, The *Sert3*-Gal4 driver target an SE1 serotonergic neuron in the SEZ (in green, GFP expression and in magenta, serotonin). **B** to **D,** SE1 neuron morphology was visualized using a membrane-bound red fluoresce protein (RFP). SerT::GFP expression (in green) changes upon co-expression of **C**, InR^CA^, or **D**, InR^DN^. The scale bar represents 10 µm. **E** to **G**, 5% sucrose intake was measured over 24 h. Expression of SerT::GFP under the control of *Sert*-Gal4 did not alter sucrose intake, but expression of **F**, InR^CA^, or **G**, InR^DN^. Combing expression with SerT::GFP restored reduced appetite when **F,** InR^CA^ was co-expressed but not when **G**, InR^DN^ was co-expressed. Data were compared using ANOVA Tukey HSD test. ** *P* < 0.01.

To test whether the activated insulin receptor indeed alters the localization of SerT, we investigated whether overexpression of the serotonin transporter in serotonergic neurons with activated insulin receptor function can restore sucrose appetite. If the morphology of the serotonergic neurons is altered, overexpression of SerT should not alter the reduced appetite for sucrose. However, if the localization of SerT is affected by the expression of the activated InR, increasing expression level of SerT could potentially bypass the effect of activated InR signaling.

We first determined whether overexpression of the serotonin transporter altered sucrose appetite (Figure 6E), which it did not. Next, we co-expressed the activated insulin receptor or blocked insulin receptor and *UAS-SerT::GFP* transgene under the control of the *Sert3*-Gal4 driver and analyzed sucrose appetite (Figure 6F and Figure G). As expected, flies with activated insulin receptor expression reduced their sucrose appetite (Figure 6D). Overexpression of SerT combined with expression of the activated insulin receptor resulted in normal sucrose appetite (Figure 6E). In contrast, blocking of insulin receptor function and co-expression of SerT did not improve appetite for sucrose (Figure 6F). Thus, the mechanisms underlying the reduced sucrose appetite induced by expression of InR^CA^ and InR^DN^ are different. The activated insulin receptor disrupts the localization of the serotonin transporter, whereas blocking the insulin receptor affects the growth of the serotonergic neurons.

In human placental cells, reduction of insulin signaling leads to retention of SerT in the endoplasmatic reticulum (ER) via an ERp44 dependent process (Kilic, 2023). To test whether an ERp44 related process could affect sucrose appetite, we analyzed the function of putative ERp44 homologs in sucrose appetite. The closet relatives of ERp44 in *Drosophila* are CG9911 with 51 % identity (Bi et al., 2021) and CG10029 with 43 % identity. Expression of RNAi transgenes of either CG9911 or CG10029 under the control of *Sert3*-Gal4 diver did not alter sucrose appetite (Figure 7A). However, we cannot rule out that knockdown of CG911 or CG 10029 is sufficient to alter sucrose appetite. Other mechanisms important for the subcellular localization of SerT depend on the coat complex II component Sec24. Mutations in Sec24 result in SerT being localized to the somatodendritic area of the neuron rather than to synaptic varicosities (Montgomery et al., 2014). To determine whether Sec24 is required for sucrose appetite, we examined the effect of Sec24_AB_ reduction in serotonergic neurons on sucrose appetite using RNAi transgenes under the control of the *Sert3*-Gal4 driver. Reduced Sec24_AB_ function reduced sucrose appetite (Figure 7B). The results are consistent with the idea that SerT localization is essential for regulating appetite-driven sucrose intake.

**Figure 7.**
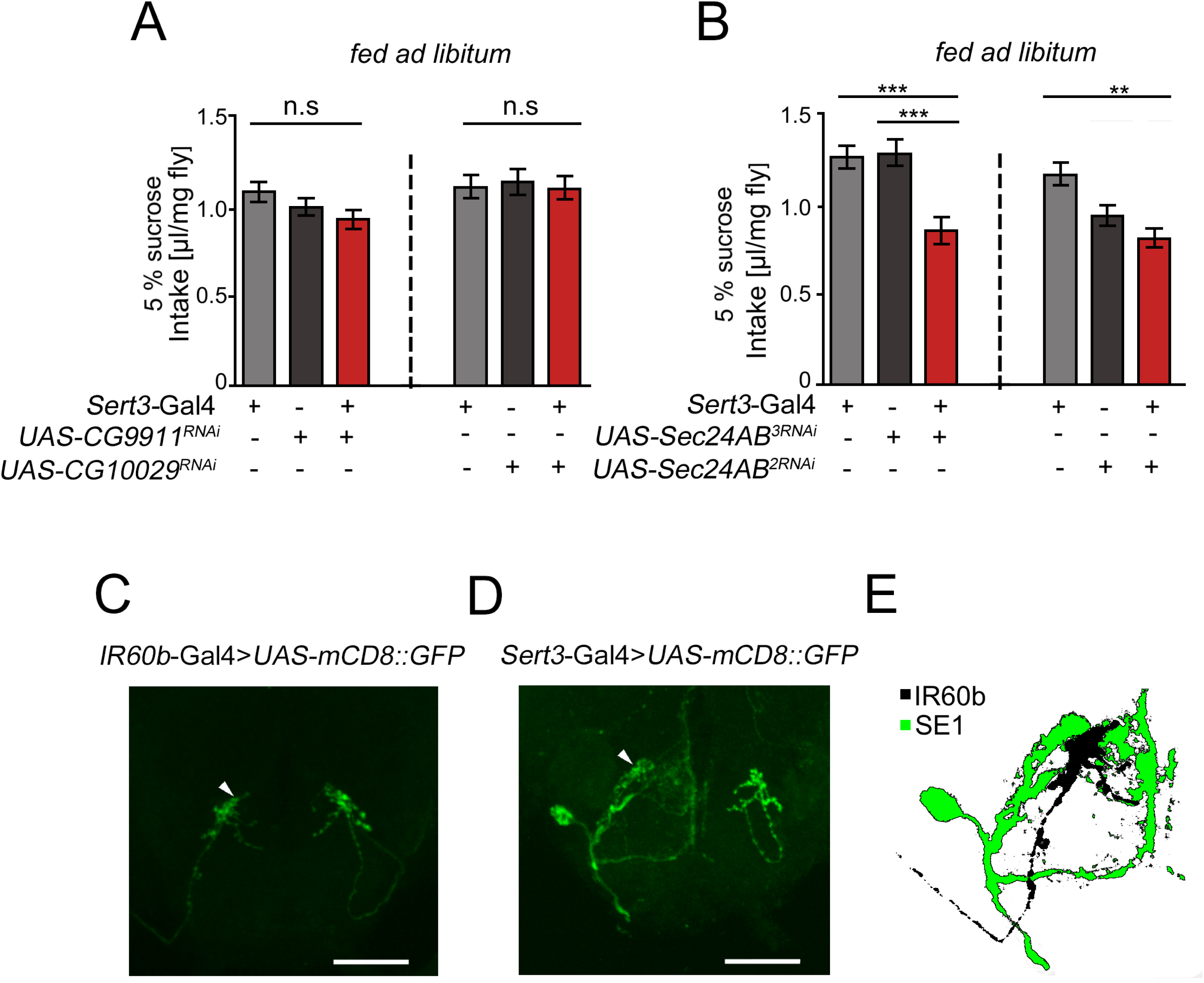
Potential pathways for regulation of sucrose appetite. **A**, Expression of *CG9911^RNAi^* or *CG10029^RNAi^* under the control of the *Sert*-Gal4 driver does not alter sucrose appetite. **B**, Reduction of Sec24AB using an RNAi transgene on the third chromosome significantly reduced sucrose appetite. Sucrose appetite of flies carrying the RNAi transgene on the second chromosome under the control of the *Sert*-Gal4 driver is significantly reduced in compared to flies carrying one copy of the Gal4 driver. Data were compared using ANOVA Tukey HSD test. ** *P* < 0.01, *** *P* < 0.001. N = 25 - 30. For raw data see Supplementary Table S1. **C**, The taste neuron IR60b and **D**, the SE1 neuron are visualized using an *UAS*-mCD8::GFP transgene. The scale bar corresponds to 10 µm. **E**, Overlay of the IR60b neuron projection with the SE1 neuron projection pattern in the SEZ.

In the SEZ, serotonin might regulate the sucrose appetite by altering the sensitivity of gustatory receptor neurons. To support this possibility, we screened the projection pattern of sucrose sensitive taste neurons (Chen & Dahanukar, 2017) (Joseph et al., 2017) in relation to the projections of the SE1 neuron. Indeed, we find that the SE1 neuron projects to the close proximity of the IR60B neuron in the SEZ (Figure 7C to Figure 7E). Neurons expressing IR60b taste receptor negatively regulate sucrose intake (Joseph et al., 2017). This observation suggests that serotonergic neurons might influence sucrose appetite by regulating taste neurons that suppress sucrose intake.

## DISCUSSION

Activation of serotonergic neurons causes both, an increase and a suppression of food intake. The level of the food intake depends on the internal energy state of the animal and the food source. Different groups of serotonergic neurons regulating the appetite for sucrose and the hunger for proteins. The serotonergic neurons that regulate the appetite for sucrose can be separated from neurons that regulate hunger-driven sucrose intake. Activation of a broader group of serotonergic neurons using the *R50H05*-Gal4 driver elicit feeding behavior in satiated flies. This suggests that among the broader set of serotonergic neurons, there are also neurons that regulate the hunger-induced food intake in satiated flies (Albin et al., 2015). We did not find, that in starved animals, activation of the same group of serotonergic neurons increases hunger for sucrose or a mixed diet, but hunger for protein enriched diet. This suggests that food intake is regulated by different mechanisms depending on the internal energy levels. This is consistent with results showing that hunger-induced sucrose uptake can be regulated by neurons upstream of Dilp--secreting neurons in the IPC (Lin et al., 2019). The regulation of Dipls is hunger-dependent (Geminard et al., 2009). Reduction of circulating hemolymph glucose can lead to a sense of hunger. The internal nutritional content of carbohydrates is directly or indirectly integrated into cell of the IPC that regulate the expression of the Dilps (Lin et al., 2019; Galikova & Klepsatel, 2023). The requirement of the serotonin receptor 5-HT1A in the IPC for food intake (Luo et al., 2014) supports that hunger-induced sucrose intake might be in part also regulated by serotonin.

Small serotonergic neurons located in the SEZ transmit information of sucrose taste to the IPC (Yao & Scott, 2022). Information about of internal glucose storage, conveyed via Dilp signaling, and external information about a sucrose source, perceived via taste, must be integrated. We identified a small group of serotonergic neurons that are regulated by the insulin receptor and reduce nutrient-specific appetite for sucrose. This group of neurons also includes the serotonergic SE1 neuron, which is located in the SEZ, responds to bitter taste and activates enteric neurons that stimulate gastric motility (Yao & Scott, 2022). In rats, increased peripheral serotonin levels reduce food intake, but do not induce satiety-related behavior (Simansky & Vaidya, 1990) making it rather unlikely that gastric motility regulates appetite-driven sucrose intake.

Alternatively, SE1 neuron function depends on the strength of stimulation. A relative “weak” sucrose stimulus leads to reduced food intake involving the IR60b neuron, and a stronger bitter stimulus increases gastric motility. The reduction of sucrose appetite correlates with a reduction of other appetite-driven behaviors such as a reduction in positive association of a food rewarded odorant or a reduced foraging behavior (Xu et al., 2016). The loss of appetite affects how the animal perceives a food reward and evaluates a food-associated odor stimulus.

Activation of a broader group of serotonergic neurons regulates nutrient-specific hunger for protein. Hunger shifts a preference for consuming sucrose to a preference for consuming yeast-containing sucrose (Ro et al., 2016). We do not detect an increase in hunger for yeast-containing sucrose, suggesting that the increase in protein hunger is not due to a shift in preference. In addition to the serotonergic neurons that regulate appetite for sucrose, there are serotonergic neurons that regulate hunger for protein. The information of sucrose appetite and protein hunger is offset against each other because hungry flies fed normal levels of food containing sucrose and yeast.

### InR and SerT regulate the appetite for sucrose

At the cellular level, at least two distinct insulin receptor-dependent functions contribute to the regulation of sucrose appetite. First, InR signaling regulates the morphology of serotonergic neurons that regulate sucrose appetite. At the cellular level, blocking insulin receptor signaling reflects a state of nutrient deprivation and can lead to reduced cellular growth (Britton et al., 2002). Blocking insulin receptor signaling in neuropeptidergic neurons reduces growth of neurite branch growth (Gu et al., 2014) and loss of insulin receptor function results in axon guidance defects of photoreceptor cells (Song et al., 2003). On a smaller scale, in mechanosensory neurons, the insulin receptor also regulates terminal synapse growth in an axon branch-specific manner (Urwyler et al., 2019). We observe that blocking InR function leads to a reduction in SE1 neuron branching in the SEZ and that blocking insulin receptor function in the central complex leads to a more diffuse expression of serotonin transporters in a region of synaptic output. We cannot distinguish whether the morphological defects of the serotonergic neurons are due to defects in axonal outgrowth and/or growth defects. However, due to the morphological changes, the SE1 neuron does not project to the correct target region. The morphological changes indirectly affect serotonin transporter function, since expression of SerT in serotonergic neurons with blocked InR function does not improve the reduced appetite for sucrose.

The second mechanism involves a more direct interaction between the insulin receptor and the serotonin transporter. Activation of the insulin receptor leads to dislocation of the serotonin transporter. The reduced sucrose uptake caused by activation of the insulin receptor can be restored by over-expression of the serotonin transporter. The function of the serotonin transporter depends on post-translational modifications (Cooper et al., 2019). Serotonin reuptake by the serotonin transporter is inhibited by phosphorylation of the serotonin transporter, increased internalization of the serotonin transporter, or reduced trafficking of serotonin transporter-containing membrane vesicles to the plasma membrane (Cooper et al., 2019). Recent studies show that in the placenta, the insulin receptor together with specific proteins of the endoplasmic reticulum regulates the trafficking of SERT to the plasma membrane (Kilic, 2023). A similar mechanism may also apply to neurons, since the knockdown of the ER protein Sec24A/B leads to a similarly reduced appetite for sucrose as the modulation of serotonin transporter function and alteration of insulin receptor signaling. Taken together, in serotonergic neurons, insulin receptor signal transduction together with the serotonin transporter enables the selective evaluation of external food sources and internal energy stores. This interaction contributes to the selective replenishment of internal energy supplies.

## STAR METHODS

### Key Resources Table

| Reagent or Resource | Source | Identifier |
| --- | --- | --- |
| **Antibodies** |  |  |
| mouse anti-nc82 (1:20) | DSHB | Cat# nc82, RRID:AB_2314866 |
| mouse anti-GFP (1:200) | Sigma Aldrich | Cat# G6539, RRID:AB_259941 |
| rabbit anti-GFP (1:1000) | Rockland | Rockland Cat# 600-401-215, RRID:AB_828167 |
| rabbit anti-SerT (1:5000) | Scholz lab |  |
| Chicken anti-RFP (1:1000) | Rockland | Cat# 600-901-379<br>RRID:AB_10704808 |
| goat anti-mouse-Cy3 (1:500) | Jackson ImmunoResearch | Cat# 115-165-146, RRID:AB_2338690 |
| goat anti-mouse-Alexa488 (1:200) | Invitrogen | Cat# A-11003, RRID:AB_141370 |
| goat anti-rabbit-Cy3 (1:1000) | Jackson ImmunoResearch | Cat# 111-165-003, RRID:AB_2338000 |
| goat anti-rabbit-Alexa488 (1:1000) | Invitrogen | Cat# A-11008, RRID:AB_143165 |
| Donkey anti-chicken-Cy3 (1:200) | Merck | Cat#6C0184<br>Product code: AP194C |
| **Chemicals, peptides and recombinant proteins** |  |  |
| All *trans*-Retinal | Sigma-Aldrich | Cat# 24899355; CAS: 116-31-4 |
| 3-octanol | Sigma-Aldrich | Cat# 93856; CAS: 589-98-0 |
| 4-methylcyclohexanol | Sigma-Aldrich | Cat# 153095; CAS: 589-91-3 |
| paraffin oil | Sigma-Aldrich | Cat# 76235; CAS: 8012-95-1 |
| red food dye | Ruth | Cat#:1.9206.1000 |
| **Experimental models: Organisms/strains** |  |  |
| *Drosophila melanogaster*: w[1118] | Scholz lab |  |
| *Drosophila melanogaster*: w[1118];IR60b-Gal4 | (Joseph et al., 2017) |  |
| *Drosophila melanogaster*: w[1118];Sert3-Gal4 | (Xu et al., 2016) |  |
| *Drosophila melanogaster*: w[1118];Tph-Gal4 | (Park et al., 2006) |  |
| *Drosophila melanogaster*: w[1118];R50H05-Gal4 | (Jenett et al., 2012) |  |
| *Drosophila melanogaster*: y[1] w[1118]; P{w[+mC]=UAS-InR.K1409A}3 (UAS-InR<sup>DN</sup>) | Bloomington Drosophila Stock center | RRID:BDSC_8253 |
| *Drosophila melanogaster*: y[1] w[1118]; P{w[+mC]=UAS-InR.A1325D}2 (UAS-InR<sup>CA</sup>) | Bloomington Drosophila Stock center | RRID:BDSC_8263 |
| *Drosophila melanogaster*: y[1] v[1]; P{y[+t7.7] v[+t1.8]=TRiP.HMJ23478}attP40 (UAS-CG10029<sup>RNAi</sup>) | Bloomington Drosophila Stock center | RRID:BDSC_61922 |
| *Drosophila melanogaster*: y[1] v[1]; P{y[+t7.7] v[+t1.8]=TRiP.JF03285}attP2 | Bloomington Drosophila Stock center | RRID:BDSC_29606 |
| (UAS-CG9911<sup>RNAi</sup>) |  |  |
| *Drosophila melanogaster*:<br>w[1118];;UAS-Sec24AB<sup>RNAi</sup> (Chr.3) | Vienna Drosophila<br>Stock Center | VDRC_44461 |
| *Drosophila melanogaster*:<br>w[1118];;UAS-Sec24AB<sup>RNAi</sup> (Chr.2) | Vienna Drosophila<br>Stock Center | VDRC_107154 |
| *Drosophila melanogaster*:<br>w[1118];;UAS-SerTDN | (Xu et al., 2016) |  |
| *Drosophila melanogaster*:<br>w[1118];;UAS-SerT::GFP | Scholz lab |  |
| *Drosophila melanogaster*: *norpA1*;<br>UAS-ChR2; UAS-ChR2 | (Classen &<br>Scholz, 2018) |  |
| **Software and algorithms** |  |  |
| Fiji/ImageJ |  | https://imagej.net/software/fiji/ |
| Excel | Microsoft<br>Corporation | https://www.microsoft.com/de-de/microsoft-365/excel?rtc=1 |
| Prism | GraphPad | https://graphpad.com/scientific-software/prism/ |
| Gimp | Das GIMP-Team | https://www.gimp.org/ |
| **Other** |  |  |
| Micro hematocrit capillaries | Brand | Cat# 749311 |
| Capillaries (5 µl) | Blaubrand | Cat# 701900 |

### Lead contact

Further information and requests for resources and reagents should be directed to and will be fulfilled by the lead contact.

### Materials Availability

All materials generated will be available upon request. All behavioral data related to figures are included in the supplement Table S1.

### Experimental model and subject details

*Drosophila melanogaster* was used for all experiments. Fly stocks are listed in the key resources table. Flies were bred and maintained on standard agar-corn meal-based food at 25 °C and 60 % relative humidity under a 12-h light-dark cycle. Flies carrying temperature sensitive transgenes were maintained at 18 °C. Experimental flies were grown in a density-controlled manner. Three- to five-day-old male flies recovered from CO_2_ sedation for 48h were used for all experiments. Flies carrying transgenes were backcrossed to the Scholz lab *w^1118^* background for five generation prior to experiments.

### Methods

#### Food intake

Food intake was measured according (Diegelmann et al., 2017). To measure how much food hungry flies feed, 20 flies were starved for 18h in vials with a humidified Whatman filter and then fed from two 5-µl capillaries containing food for 3h. To measure appetite, 8 non-starved flies had *ad libitum* access to food from four 5-µl capillaries for 24h. The food solutions were colored with 2% red food dye (Ruth GmbH). The amount of food was measured using a caliper. To account for evaporation, the evaporation rate of three assays without flies were determined. Food intake was normalized to the evaporation rate, the number of flies and body weight. The body weight was determined from at least 5 populations of 100 flies per genotype.

To analyze the role of acute activation of serotonergic neurons in food intake, the capillary feeder assay was combined with an optogenetic set up described in detail in (Schneider et al., 2012). Briefly, a blue LED (465 – 485 nm; Cree; Germany) with light intensity of 1800 lux was placed approximately 36 mm above the capillary feeder assay. The light was pulsed with a sequence of 2s of 40Hz, 16s of 8Hz and 2s without light. The control group was reared with standard food containing the solvent and the experimental group with standard food containing 300µl of 250mM all-trans-retinal dissolved in 100% ethanol. The flies were kept in the dark. To recover from CO_2_ sedation, flies were kept in vials containing vehicle or all-trans-retinal for 2d prior to experiments.

#### Learning and memory

Associative olfactory short-term learning and memory tests were performed according to (Tully & Quinn, 1985) with modification of (Schwaerzel et al., 2003). For appetitive short-term memory, flies were starved for 16 to 20h prior to the experiment. For training, 100 3- to 5-day-old male flies were exposed to the first odorant without reinforcer (conditioned stimulus minus) for 2 min, followed by an exposure to a second odorant paired with 2M sucrose (conditioned stimulus plus). For aversive short-term memory, flies were not starved. Flies were first exposed to the odorant paired with electric shock (12 1.3s pulses at 90V spaced in 5s apart), followed by exposure to the second odorant. After training, flies chose between the two odorants for 2min. The performance index (PI_1/2_) was calculated as the PI_1/2_ = (#_CS+_ - #_CS-_)/ (#_CS+_ + #_CS-_). To exclude non-associative effects, each experiment consisted of two independent reciprocal groups of flies. The learning index was calculated from the average scores of two PIs as follows LI = (PI_1_ + PI_2_)/2). Training and testing were conducted in a T-maze at 25 °C and 80% relative humidity. The odorants used were 3-octanol (1:80 diluted in paraffin oil) and 4-methylcyclohexanol (1:100 diluted in paraffin oil).

#### Statistics

The Shapiro-Wilk Test with α = 0.05 was used to determine whether the data were normally distributed. Two groups of normally distributed data were compared using unpaired Student’s t-test assuming equal variance and more than two groups with one-way ANOVA, with Tukey’s post-hoc HSD test (*P\** < 0.05; *P\*\** < 0.01; *P\*\*\** < 0.001). The statistics were performed using https://www.statskingdom.com/.

#### Immunohistochemistry

Fly brains were dissected in ice-cold *Drosophila* Ringer and transferred to phosphate-buffered saline (PBS, pH 7.4). Brains were fixed with 3.7 % formaldehyde for 25 min. Tissue was washed at least three times for 15 min with PBT (0.5 % Triton X-100). Brains were blocked with 5 % fetal calf serum in PBT for at least 1.5 h. A primary antibody solution was then added and brains were incubated overnight at 4 °C. Brains were washed with 0.5 % PBST and incubated with the secondary antibody for at least 3 h at room temperature or overnight at 4 °C. brains were then washed again three times with 0.5 % PBST and embedded in VectaShield^©^. Confocal images were acquired with an Olympus FV1000MPE or Leica SP8 laser scanning microscope and analyzed with Fiji ImageJ version 1.53c-q.

## ACKNOWLEDGEMENTS

We thank the Bloomington *Drosophila* Stock Center for providing *Drosophila* fly lines. H.S. was supported by TR 1340, DFG-Scho656_10-1, DFG-Scho656_14-1 and the Hetzler award, K.D. and H.S. RTG1960.

## AUTHOR CONTRIBUTIONS

H.S. initiated the project. M.G., K.D., J.H. and H.S. designed the experiments. M.G., K.D. and J.H performed the experiments and, together with H.S., analyzed the data. M.G. and K.D. wrote drafts of the manuscript, and H.S. wrote the manuscript.

## DECLARARATION OF INTERESTS

The authors declare no competing interests.

---

**Supplementary Table S1. Raw data related to Figure 1 to Figure 5 and Figure 7**. The supplementary Table S1 contains the raw behavioral data used to generate the figures.

## REFERENCES

Albin, S. D., Kaun, K. R., Knapp, J. M., Chung, P., Heberlein, U., & Simpson, J. H. (2015). A Subset of Serotonergic Neurons Evokes Hunger in Adult Drosophila. Curr Biol, 25(18), 2435–2440.

Berthoud, H. R., & Morrison, C. (2008). The brain, appetite, and obesity. Annu Rev Psychol, 59, 55–92.

Bi, Y., Chang, Y., Liu, Q., Mao, Y., Zhai, K., Zhou, Y., et al. (2021). ERp44/CG9911 promotes fat storage in Drosophila adipocytes by regulating ER Ca(2+) homeostasis. Aging (Albany NY), 13(11), 15013–15031.

Britton, J. S., Lockwood, W. K., Li, L., Cohen, S. M., & Edgar, B. A. (2002). Drosophila’s insulin/PI3-kinase pathway coordinates cellular metabolism with nutritional conditions. Dev Cell, 2(2), 239–249.

Chatterjee, N., & Perrimon, N. (2021a). What fuels the fly: Energy metabolism in Drosophila and its application to the study of obesity and diabetes. Sci Adv, 7(24).

Chatterjee, N., & Perrimon, N. (2021b). What fuels the fly: energy metabolism in *Drosophila* and its application to the study of obesity and diabetes. Sci. Adv., 7(24), eabg4336.

Chen, Y. D., & Dahanukar, A. (2017). Molecular and Cellular Organization of Taste Neurons in Adult Drosophila Pharynx. Cell Rep, 21(10), 2978–2991.

Chowanski, S., Walkowiak-Nowicka, K., Winkiel, M., Marciniak, P., Urbanski, A., & Pacholska-Bogalska, J. (2021). Insulin-Like Peptides and Cross-Talk With Other Factors in the Regulation of Insect Metabolism. Front Physiol, 12, 701203.

Classen, G., & Scholz, H. (2018). Octopamine Shifts the Behavioral Response From Indecision to Approach or Aversion in Drosophila melanogaster. Front Behav Neurosci, 12, 131.

Cooper, A., Woulfe, D., & Kilic, F. (2019). Post-translational modifications of serotonin transporter. Pharmacol Res, 140, 7–13.

Demontis, F., & Perrimon, N. (2009). Integration of Insulin receptor/Foxo signaling and dMyc activity during muscle growth regulates body size in Drosophila. Development, 136(6), 983–993.

Diegelmann, S., Jansen, A., Jois, S., Kastenholz, K., Velo Escarcena, L., Strudthoff, N., & Scholz, H. (2017). The CApillary FEeder Assay Measures Food Intake in Drosophila melanogaster. J Vis Exp(121).

Galikova, M., & Klepsatel, P. (2023). Endocrine control of glycogen and triacylglycerol breakdown in the fly model. Semin Cell Dev Biol, 138, 104–116.

Geminard, C., Rulifson, E. J., & Leopold, P. (2009). Remote control of insulin secretion by fat cells in Drosophila. Cell Metab, 10(3), 199–207.

Giang, T., Ritze, Y., Rauchfuss, S., Ogueta, M., & Scholz, H. (2011). The serotonin transporter expression in Drosophila melanogaster. J Neurogenet, 25(1-2), 17–26.

Gu, T., Zhao, T., & Hewes, R. S. (2014). Insulin signaling regulates neurite growth during metamorphic neuronal remodeling. Biol Open, 3(1), 81–93.

Hainer, V., Kabrnova, K., Aldhoon, B., Kunesova, M., & Wagenknecht, M. (2006). Serotonin and norepinephrine reuptake inhibition and eating behavior. Ann N Y Acad Sci, 1083, 252–269.

Halford, J. C., Harrold, J. A., Boyland, E. J., Lawton, C. L., & Blundell, J. E. (2007). Serotonergic drugs : effects on appetite expression and use for the treatment of obesity. Drugs, 67(1), 27–55.

Hanesch, U., Fischbach, K. F., & Heisenberg, M. (1989). Neuronal Architecture of the Central Complex in Drosophila-Melanogaster. Cell and Tissue Research, 257(2), 343–366.

He, J., Hommen, F., Lauer, N., Balmert, S., & Scholz, H. (2020). Serotonin transporter dependent modulation of food-seeking behavior. PLoS One, 15(1), e0227554.

Inoue, Y. H., Katsube, H., & Hinami, Y. (2018). *Drosophila* models to investigate insulin action and mechanisms underlying human diabetes mellitus. Adv. Exp. Med. Biol., 1076, 235–256.

Ja, W. W., Carvalho, G. B., Mak, E. M., de la Rosa, N. N., Fang, A. Y., Liong, J. C., et al. (2007). Prandiology of Drosophila and the CAFE assay. Proc Natl Acad Sci U S A, 104(20), 8253–8256.

Jenett, A., Rubin, G. M., Ngo, T. T., Shepherd, D., Murphy, C., Dionne, H., et al. (2012). A GAL4-driver line resource for Drosophila neurobiology. Cell Rep, 2(4), 991–1001.

Joseph, R. M., Sun, J. S., Tam, E., & Carlson, J. R. (2017). A receptor and neuron that activate a circuit limiting sucrose consumption. Elife, 6.

Kilic, F. (2023). The nature of the binding between insulin receptor and serotonin transporter in placenta (review). Placenta, 133, 40–44.

Knapp, E. M., Kaiser, A., Arnold, R. C., Sampson, M. M., Ruppert, M., Xu, L., et al. (2022). Mutation of the Drosophila melanogaster serotonin transporter dSERT impacts sleep, courtship, and feeding behaviors. PLoS Genet, 18(11), e1010289.

Ko, K. I., Root, C. M., Lindsay, S. A., Zaninovich, O. A., Shepherd, A. K., Wasserman, S. A., et al. (2015). Starvation promotes concerted modulation of appetitive olfactory behavior via parallel neuromodulatory circuits. Elife, 4.

Li, C. R., Guo, D., & Pick, L. (2013). Independent signaling by Drosophila insulin receptor for axon guidance and growth. Front Physiol, 4, 385.

Lin, S., Senapati, B., & Tsao, C. H. (2019). Neural basis of hunger-driven behaviour in Drosophila. Open Biol, 9(3), 180259.

Montgomery, T. R., Steinkellner, T., Sucic, S., Koban, F., Schuchner, S., Ogris, E., et al. (2014). Axonal targeting of the serotonin transporter in cultured rat dorsal raphe neurons is specified by SEC24C-dependent export from the endoplasmic reticulum. J Neurosci, 34(18), 6344–6351.

Nagel, G., Szellas, T., Huhn, W., Kateriya, S., Adeishvili, N., Berthold, P., et al. (2003). Channelrhodopsin-2, a directly light-gated cation-selective membrane channel. Proc Natl Acad Sci U S A, 100(24), 13940–13945.

Omoto, J. J., Nguyen, B. M., Kandimalla, P., Lovick, J. K., Donlea, J. M., & Hartenstein, V. (2018). Neuronal Constituents and Putative Interactions Within the Drosophila Ellipsoid Body Neuropil. Front Neural Circuits, 12, 103.

Park, J., Lee, S. B., Lee, S., Kim, Y., Song, S., Kim, S., et al. (2006). Mitochondrial dysfunction in Drosophila PINK1 mutants is complemented by parkin. Nature, 441(7097), 1157–1161.

Pooryasin, A., & Fiala, A. (2015). Identified Serotonin-Releasing Neurons Induce Behavioral Quiescence and Suppress Mating in Drosophila. J Neurosci, 35(37), 12792–12812.

Ro, J., Pak, G., Malec, P. A., Lyu, Y., Allison, D. B., Kennedy, R. T., & Pletcher, S. D. (2016). Serotonin signaling mediates protein valuation and aging. Elife, 5.

Rudnick, G., & Sandtner, W. (2019). Serotonin transport in the 21st century. J Gen Physiol, 151(11), 1248–1264.

Saltiel, A. R., & Kahn, C. R. (2001). Insulin signalling and the regulation of glucose and lipid metabolism. Nature, 414(6865), 799–806.

Schwaerzel, M., Monastirioti, M., Scholz, H., Friggi-Grelin, F., Birman, S., & Heisenberg, M. (2003). Dopamine and octopamine differentiate between aversive and appetitive olfactory memories in Drosophila. J Neurosci, 23(33), 10495–10502.

Semaniuk, U., Gospodaryov, D., Mishchanyn, K., Storey, K., & Lushchak, O. (2021). Drosophila insulin-like peptides regulate concentration-dependent changes of appetite to different carbohydrates. Zoology (Jena), 146, 125927.

Semaniuk, U. V., Gospodaryov, D. V., Feden’ko, K. M., Yurkevych, I. S., Vaiserman, A. M., Storey, K. B., et al. (2018). Insulin-Like Peptides Regulate Feeding Preference and Metabolism in Drosophila. Front Physiol, 9, 1083.

Simansky, K. J., & Vaidya, A. H. (1990). Behavioral mechanisms for the anorectic action of the serotonin (5-HT) uptake inhibitor sertraline in rats: comparison with directly acting 5-HT agonists. Brain Res Bull, 25(6), 953–960.

Song, J., Wu, L., Chen, Z., Kohanski, R. A., & Pick, L. (2003). Axons guided by insulin receptor in Drosophila visual system. Science, 300(5618), 502–505.

Sudhakar, S. R., Pathak, H., Rehman, N., Fernandes, J., Vishnu, S., & Varghese, J. (2020). Insulin signalling elicits hunger-induced feeding in Drosophila. Dev Biol, 459(2), 87–99.

Tierney, A. J. (2020). Feeding, hunger, satiety and serotonin in invertebrates. Proc Biol Sci, 287(1932), 20201386.

Tully, T., & Quinn, W. G. (1985). Classical conditioning and retention in normal and mutant Drosophila melanogaster. J Comp Physiol A, 157(2), 263–277.

Urwyler, O., Izadifar, A., Vandenbogaerde, S., Sachse, S., Misbaer, A., & Schmucker, D. (2019). Branch-restricted localization of phosphatase Prl-1 specifies axonal synaptogenesis domains. Science, 364(6439).

Voigt, J. P., & Fink, H. (2015). Serotonin controlling feeding and satiety. Behav Brain Res, 277, 14–31.

Xu, L., He, J., Kaiser, A., Graber, N., Schlager, L., Ritze, Y., & Scholz, H. (2016). A Single Pair of Serotonergic Neurons Counteracts Serotonergic Inhibition of Ethanol Attraction in Drosophila. PLoS One, 11(12), e0167518.

Yabut, J. M., Crane, J. D., Green, A. E., Keating, D. J., Khan, W. I., & Steinberg, G. R. (2019). Emerging Roles for Serotonin in Regulating Metabolism: New Implications for an Ancient Molecule. Endocr Rev, 40(4), 1092–1107.

Yao, Z., & Scott, K. (2022). Serotonergic neurons translate taste detection into internal nutrient regulation. Neuron, 110(6), 1036–1050 e1037.

